# Improved Genome Packaging Efficiency of AAV Vectors Using Rep Hybrids

**DOI:** 10.1101/2021.05.10.443533

**Authors:** Mario Mietzsch, Courtnee Eddington, Ariana Jose, Jane Hsi, Paul Chipman, Tom Henley, Modassir Choudhry, Robert McKenna, Mavis Agbandje-McKenna

**Affiliations:** Department of Biochemistry and Molecular Biology, Center for Structural Biology, McKnight Brain Institute, College of Medicine, University of Florida, Gainesville, FL 32610, USA; Intima Bioscience, New York, USA

**Keywords:** Adeno-associated virus, Rep protein, capsid, genome packaging, cryo-electron microscopy, hybrid, empty and full capsids

## Abstract

Recombinant Adeno-associated viruses (rAAVs) are one of the most commonly used vectors for a variety of gene therapy applications. In the last two decades research focused primarily on the characterization and isolation of new *cap* genes resulting in hundreds of natural and engineered AAV capsid variants while the *rep* gene, the other major AAV open reading frame, has been less studied. This is due to the fact that the *rep* gene from AAV serotype 2 (AAV2) enables the ssDNA packaging of recombinant genomes into most AAV serotype and engineered capsids. However, a major byproduct of all vector productions is empty AAV capsids, lacking the encapsidated vector genome, especially for non-AAV2 vectors. Despite the packaging process being considered the rate-limiting step for rAAV production, none of the *rep* genes from the other AAV serotypes have been characterized for their packaging efficiency. Thus, in this study AAV2 *rep* was replaced with the *rep* gene of a select number of AAV serotypes. However, this led to a lowering of capsid protein expression, relative to the standard AAV2-*rep* system. In further experiments the 3’end of the AAV2 *rep* gene was reintroduced to promote increased capsid expression and a series of chimeras between the different AAV Rep proteins were generated and characterized for their vector genome packaging ability. The utilization of these novel Rep hybrids increased the percentage of genome containing (full) capsids ~2-4-fold for all of the non-AAV2 serotypes tested. Thus, these Rep chimeras could revolutionize rAAV production.

**Importance:** A major byproduct of all Adeno-associated virus (AAV) vector production systems are “empty” capsids, void of the desired therapeutic gene, and thus do not provide any curative benefit for the treatment of the targeted disease. In fact, empty capsids can potentially elicit additional immune responses *in vivo* gene therapies if not removed by additional purification steps. Thus, there is a need to increase the genome packaging efficiency and reduce the number of empty capsids from AAV biologics. The novel Rep hybrids from different AAV serotypes described in this study are capable of reducing the percentage of empty capsids in all tested AAV serotypes and improve overall yields of genome-containing AAV capsids at the same time. They can likely be integrated easily into existing AAV manufacturing protocols to optimize the production of the generated AAV gene therapy products.

## Introduction

Adeno-associated viruses (AAVs) are not associated with any pathogenic effects but are widely studied as they have been developed into one of the most promising gene therapy vectors for *in vivo* gene therapy for a wide variety of monogenetic diseases (1). To date, three AAV-vector-mediated gene therapies have gained approval for commercialization: Glybera, an AAV1 vector for the treatment of lipoprotein lipase deficiency (2); Luxturna, an AAV2 vector for the treatment of Leber’s congenital amaurosis (3); and Zolgensma, an AAV9 vector for the treatment of spinal muscular atrophy type 1 (4). Common to all natural and engineered AAVs are their small, non-enveloped icosahedral capsids, of ~260 Å diameter, that contain a linear single-stranded DNA (ssDNA) genome (5). The wild-type AAVs have a genome size of approximately 4.7 kb (6). Both ends of the genome contain identical inverted terminal repeats (ITRs) of ~150 nucleotides (nts) that form T-shaped hairpin secondary structures and are important for genome replication and packaging (7). Between these ITRs, two open reading frames (ORFs) encode a series of replication (Rep) and virus proteins (VP) (8), as well as two smaller accessory proteins: the assembly activating protein (AAP) and the membrane-associated accessory protein (MAAP) (9, 10). For recombinant AAV vectors, these viral ORFs are replaced with an approximately similar-sized therapeutic gene of interest and only the flanking *cis*-active ITRs are retained to allow packaging of the recombinant genomes into the capsids. Nevertheless, the Rep and VP proteins are also needed for vector manufacturing but can be supplied in *trans* (11, 12).

The Rep proteins play critical roles in the viral replication cycle as well as for vector production (8). The four Rep proteins, Rep78, Rep68, Rep52, and Rep40, are situated in the same ORF and are translated from transcripts generated by the p5 and p19 promoter (13). The larger Rep78/68 proteins are extended by ~224 amino acids at their N-terminus in comparison to Rep52/40 and the C-terminus of Rep68/40 differs from those of Rep78/52 due to differential splicing (13). All Rep proteins share a central 305 amino acid stretch that contains a helicase/ATPase domain (14) and nuclear localization signals (15). The N-terminus of Rep78/68 contains a DNA binding and endonuclease domain (16) whereas the C-terminus of Rep78/52 is suggested to contain a zinc finger domain (17). The large Rep proteins Rep78/68 are indispensable for genome replication and were shown to bind to a specific DNA sequence also present in the ITRs. The small Rep proteins Rep52/40 are essential for packaging of the genome into the capsids (18) formed by VP1, VP2, and VP3 that are encoded by the second ORF.

Similar to the Rep proteins the VPs are located in one ORF and share a common C-terminus. They are generated from transcripts of the p40 promoter by differential splicing and by the utilization of alternate start codons. VP1 is the largest capsid protein with approximately 735 amino acids and is encoded within the entire ORF. VP2 and VP3 are truncated at their N-termini relative to VP1 with ~598 and 533 amino acids, respectively. The VPs are expressed and incorporated into the capsid in an approximate ratio of 1:1:10 (VP1:VP2:VP3) (19). The capsid assembly is assisted by AAP which is encoded within the *cap* gene but situated in an alternate reading frame (9). Similarly, the second accessory protein MAAP is also encoded in an alternate reading frame within the *cap* gene and was suggested to have a role in capsid egress (10).

Packaging of the wild-type or recombinant vector genomes occurs in the nucleus into preformed “empty” VP capsids (20). Following replication of the genomes by the large Rep proteins, the generated ssDNA genomes are thought to be translocated into the capsid in 3’ to 5’ direction by the helicase/ATPase domain of the small Rep proteins (18). The translocation of the ssDNA into the preformed capsid is proposed to occur through a channel at the 5-fold symmetry axis of the AAV capsids (5). Since genome packaging is considered to be the rate limiting step a significant number of capsids go unpacked and remain empty. For AAV-mediated gene therapy these empty capsids are not desired as they do not provide any therapeutic benefit and may induce unwanted immune responses.

For the AAVs 13 primate AAV serotypes and numerous genotypes or engineered capsids have been described (21). The Rep and Capsid proteins of the AAV serotypes vary in amino acid sequence identity between ~52-99% (Table 1). This wide variety of available capsids is utilized in AAV-mediated gene therapy to package the transgene of choice into the capsid with the desired characteristics and tissue tropism. However, for the vast majority of manufactured AAV vectors the ITRs and *rep* gene of serotype 2 (AAV2) are utilized and only the *cap* gene is changed resulting in pseudotyped AAVs (22). While the Rep proteins of AAV2 are able to package vector genomes in all AAV serotype capsids their packaging efficiency is lower for some capsids compared to AAV2 (23). As a result, AAV vector preparations with non-AAV2 capsids contain a higher percentage of empty capsids.

**Table 1:** Comparison of the VP1 and Rep78 amino acid sequence identity of the AAV serotypes

|  | AAV1 | AAV2 | AAV3 | AAV4 | AAV5 | AAV6 | AAV7 | AAV8 | AAV9 | AAV10 | AAV11 | AAV12 | AAV13 | VP1<br>amino<br>acid<br>sequence<br>identity<br>[%] |
| --- | --- | --- | --- | --- | --- | --- | --- | --- | --- | --- | --- | --- | --- | --- |
| AAV1 |  | 83 | 87 | 63 | 58 | 99 | 85 | 84 | 82 | 85 | 66 | 60 | 87 |  |
| AAV2 | 87 |  | 88 | 60 | 57 | 83 | 82 | 83 | 82 | 84 | 63 | 60 | 88 |  |
| AAV3 | 89 | 89 |  | 63 | 58 | 87 | 85 | 86 | 84 | 86 | 65 | 61 | 94 |  |
| AAV4 | 90 | 90 | 93 |  | 53 | 63 | 63 | 63 | 62 | 63 | 81 | 78 | 65 |  |
| AAV5 | 58 | 58 | 58 | 58 |  | 58 | 58 | 58 | 57 | 57 | 53 | 52 | 58 |  |
| AAV6 | 99 | 87 | 89 | 89 | 58 |  | 85 | 84 | 82 | 85 | 66 | 60 | 87 |  |
| AAV7 | 98 | 88 | 90 | 90 | 58 | 98 |  | 88 | 82 | 88 | 67 | 62 | 85 |  |
| AAV8 | 95 | 85 | 87 | 88 | 57 | 95 | 96 |  | 85 | 93 | 65 | 62 | 85 |  |
| AAV9 | N/A | N/A | N/A | N/A | N/A | N/A | N/A | N/A |  | 86 | 64 | 60 | 84 |  |
| AAV10 | 97 | 88 | 89 | 90 | 58 | 97 | 97 | 95 | N/A |  | 66 | 61 | 86 |  |
| AAV11 | 97 | 88 | 89 | 89 | 58 | 97 | 97 | 95 | N/A | 100 |  | 84 | 65 |  |
| AAV12 | 87 | 88 | 89 | 88 | 58 | 87 | 88 | 85 | N/A | 88 | 88 |  | 60 |  |
| AAV13 | 89 | 90 | 93 | 99 | 58 | 89 | 90 | 88 | N/A | 89 | 89 | 88 |  |  |
| Rep78 amino acid sequence identity [%] |  |  |  |  |  |  |  |  |  |  |  |  |  |  |

Thus, in order to characterize the utilization of other *rep* genes in AAV vector productions, in this study the AAV2 *rep* gene was replaced with the *rep* gene of the same AAV serotype as the *cap* gene for a selection of different AAV serotypes. However, this substitution led to an overall lower VP expression for the analyzed AAV serotypes relative to the standard AAV2-*rep* system. Reverting the 3’ end of the *rep* gene back to AAV2 as well as corrections in the DNA-binding domain of AAV8 Rep restored high capsid expression. In addition, a higher packaging efficiency of vector genomes was observed. In a series of chimeras, primarily the DNA binding and endonuclease domain of the AAV1 and AAV8 Rep was shown to be responsible for improved vector genome packaging for AAV1 and AAV8 vectors. These observations were confirmed by ELISA, qPCR, alkaline gel electrophoresis, and cryo-electron microscopy (cryo-EM) of affinity-purified AAV vector preparations and show that the new Rep hybrids can increase the number of genome-containing capsids 2 to 4-fold in the case of AAV1 and AAV8 when compared to AAV2-*rep* produced vectors. Furthermore, utilization of these hybrids for other AAVs such as AAV6, AAV9, and AAVrh.10 also show higher genome packaging efficiency. These observations indicate that the utilization of these Rep chimeras could revolutionize AAV vector production.

## Results and Discussion

### Substitution of the standard AAV2 rep gene

In the last two decades a lot of research has focused on the characterization and isolation of new *cap* genes resulting in hundreds of natural and engineered AAV capsid variants (21, 24–27). At the same time, the *rep* gene has been largely ignored due to the fact that the Rep proteins of AAV2 also package recombinant vector genomes into almost all AAV serotypes or variant capsids. A major byproduct of all vector productions is empty AAV capsids, with even lower packaging efficiencies for non-AAV2 vectors (23). Compared to AAV2, the amino acid sequence identity of VP1 ranges from 82-88% for most AAV serotype capsids, with the exception of AAV4/5/11/12 (Table 1). For the foremost AAV serotypes, the amino acid sequence identity of Rep78 shows a similar range of 85-90% indicating a potential co-evolution of the Rep proteins with the capsid VPs, potentially due to optimal interaction of Rep proteins with the matched serotype capsids during genome packaging.

Thus, to understand the impact for the utilization of different AAV Rep proteins for AAV vector production, the AAV2 *rep* gene was replaced in the producer plasmids with the *rep* gene of the AAV serotype identical to the *cap* gene for the serotypes AAV1, 6, and 8 (Figure 1A). When the expression of the capsid proteins was analyzed all of new constructs demonstrated low to no VP expression (in the case of AAV8) compared to the standard AAV2 *rep* constructs (Figure 1B). The majority of AAV producer plasmids utilize AAV2 *rep* genes with the Rep78 start codon changed to an ACG, to lower the expression of the large Rep proteins in favor of the small Rep proteins (28). This strategy was shown to improve the overall AAV yield (29). Thus, ATG and ACG constructs for each producer plasmid were generated. While the Rep78 start codon had no significant effect on VP expression for any construct, the ACG start codon reduced Rep78/68 expression in all constructs as anticipated (Figure 1B) but in contrast to previous observations the overall yield of AAV vectors was not increased when using the ACG-Rep2 constructs (Figure 1C). In fact, the yield of AAV1 and AAV6 vectors using the AAV1 and AAV6 *rep* constructs showed higher yields with the Rep78 having an ATG start codon. Another observation for the AAV1 and AAV6 *rep* constructs was their lower expression levels of spliced Rep proteins, Rep68 and Rep40, compared to the AAV2 *rep* constructs. The similar behavior of the AAV1 and 6 constructs was expected as their *rep* genes differ only in 36 nucleotides resulting in only four amino acid differences or 99% sequence identity (Table 1). This level of similarity between AAV1 and 6 is comparable to VP1 with only six amino acid differences between the AAV serotypes.

**Figure 1:**
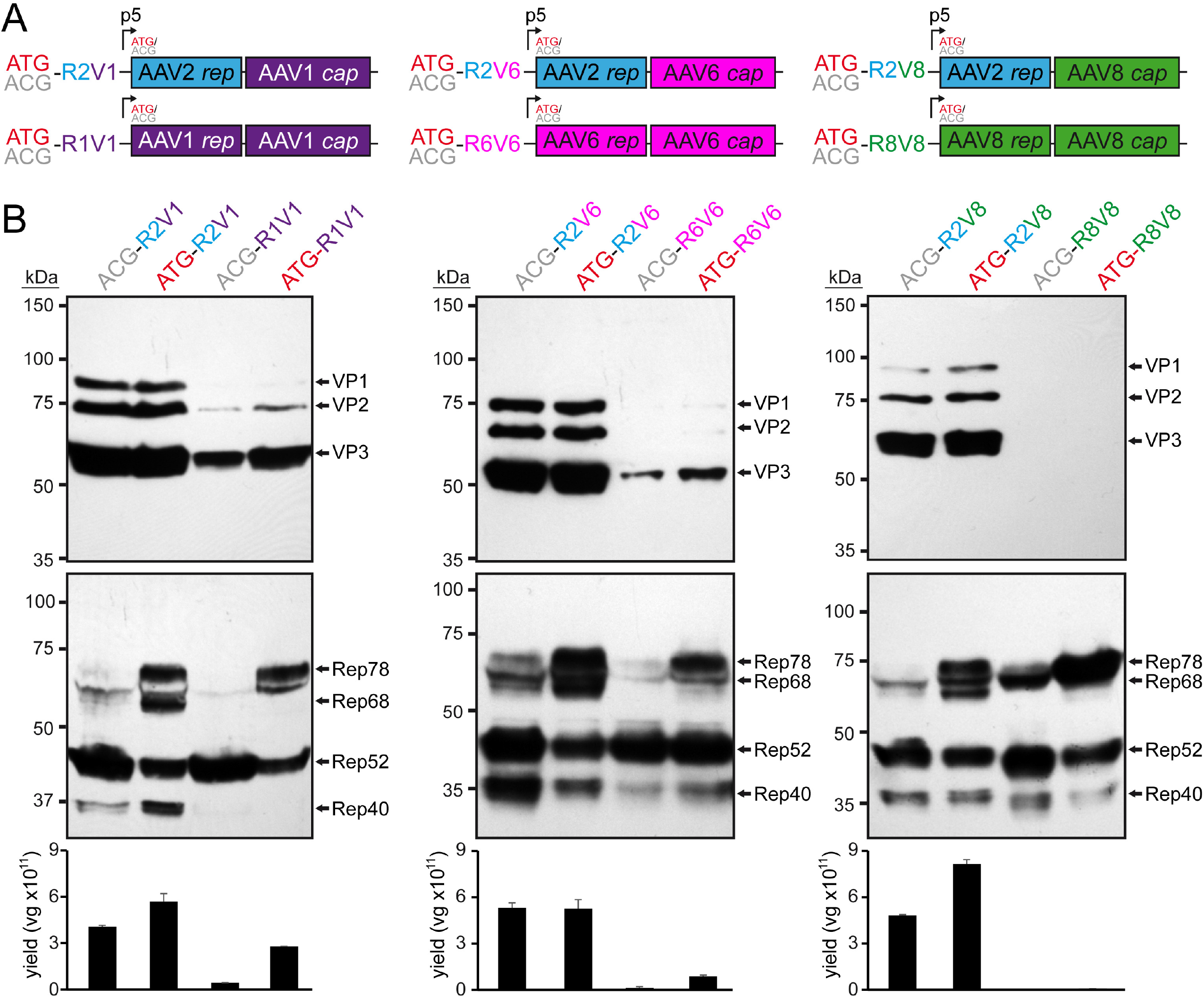
Substitution of the AAV2 *rep* gene. **A)** Overview of the utilized *rep*-*cap* constructs. AAV genes derived from AAV1 are colored in purple, AAV2 in blue, AAV6 in pink, and AAV8 in green. The construct name indicates the utilized Rep78 start codon (ATG or ACG) as well as the origin of the *rep* (R) and *cap* (V) gene (e.g. R2V1: *rep* gene from AAV2 and *cap* gene from AAV1). **B)** Analysis of Rep and VP expression by Western-blot following transfection of the constructs in HEK293 cells. Top blots were probed with MAb B1 and lower blots with MAb 1F. The individual VPs and Rep proteins are indicated.

### Reintroduction of AAV2 sequences at the 3’ end of AAV1 rep gene rescues VP expression

The low expression of the capsid proteins prevents efficient AAV vector production. As the VPs are translated from transcripts generated from the p40 promoter, which is located in the 3’ end of the *rep* gene, there was the possibility that the AAV1, 6, or 8 p40 promoter does not have same activity as the AAV2 p40 promoter. Alternatively, the observed reduced expression of Rep68/40 (Figure 1B) could indicate a disruption of mRNA splicing. The capsid proteins have been reported to be primarily translated from spliced p40 transcripts and mutations of the splice donor site resulted in a strong reduction of VP expression (30, 31). However, the splice donor consensus sequence of AAV1, 6, and 8 is identical to AAV2 (31). In order to test the first possibility, AAV p40-promoter luciferase reporter constructs were generated for AAV1, 2, and 8 and their luciferase expression analyzed (Figure 2A). Indeed, the AAV1 p40-promoter (AAV6 differs only by a single nucleotide) showed only about a quarter of the expression compared to AAV2. However, the AAV8 p40-promoter showed ~60% of the AAV2 p40-mediated expression (Figure 2A) even though VP expression was lower for the AAV8 construct compared to AAV1 (Figure 1B). Thus, there is the possibility of overlapping detrimental effects.

**Figure 2:**
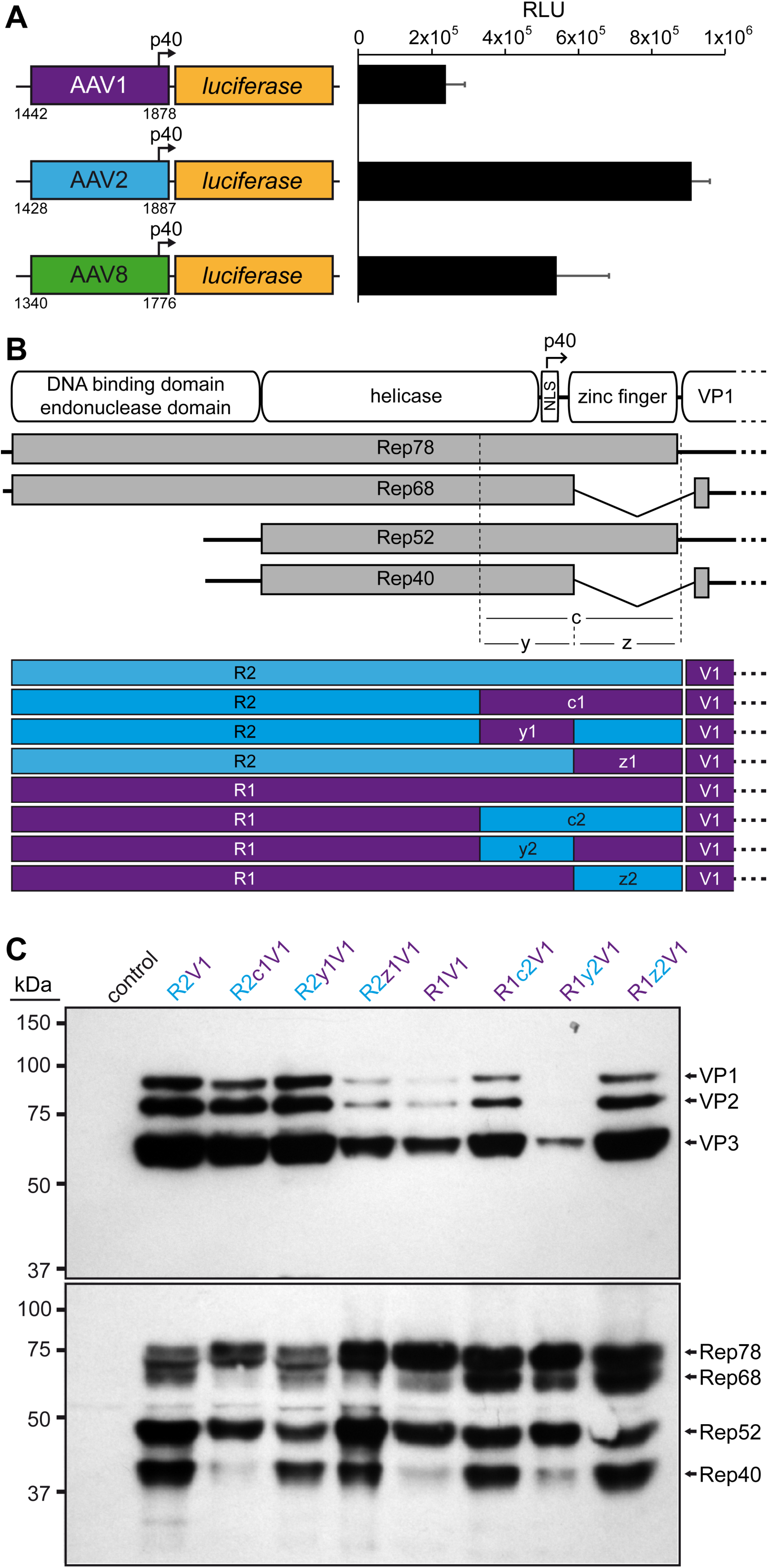
Role of the 3’ end of the rep gene in VP expression. **A)** Analysis of the p40 promoter activity from AAV1, AAV2, and AAV8 driving luciferase expression in HEK293 cells 24 h post transfection. The nucleotide positions of the fragments used are shown below the constructs based on the accession numbers: AF063497 (AAV1), AF043303 (AAV2), and AF513852 (AAV8). RLU: relative light units. **B)** Schematic depiction of the Rep protein with its main domains. The approximate position of the p40 promoter in the *rep* gene is indicated. Below the p5 and p19 transcripts are shown with the translated regions for Rep78, Rep68, Rep52, and Rep40. The C-terminal (c) ~250 aa has been subdivided into a N-terminal y-region and C-terminal z-region. The construct generated with all permutations between AAV1 and AAV2 involving these regions are depicted. Rep gene fragments derived from AAV1 are colored in purple and AAV2 in blue. **C)** Western-blot analysis to determine Rep and VP expression of the constructs. Top blots were probed with MAb B1 and lower blots with MAb 1F. The individual VPs and Rep proteins are indicated.

In order to rescue VP expression from the AAV1 *rep* constructs a series of swaps of the 3’ end of the *rep* gene (or the C-terminus on protein level) for AAV1 and 2 were generated (Figure 2B). These constructs were analyzed for their expression of the Rep and capsid proteins. As seen before the VP expression of the construct with the entire AAV1 rep (R1V1) was reduced and less Rep68/40 was observed compared to the standard AAV2 rep construct (R2V1) (Figure 2C). Substitution of the 3’ end of AAV1 *rep* to AAV2 (R2c1V2) resulted only in a minor reduction of VP expression but showed less of Rep68/40. Vice versa, the introduction of 3’ end of AAV2 *rep* in AAV1 increased VP expression and showed higher expression of the spliced Rep68/40 proteins (Figure 2C). To further narrow down the region responsible for these effects the 3’ end of the rep gene was divided into two parts. The first (y) region contains the p40-pomoter and the splice donor site whereas the second (z) region comprises the sequence encoding the zinc-finger domain (Figure 2B). Swapping the y region to the respective sequences of the other AAV serotype resulted in little to no differences of the VP and Rep expression when compared to the constructs with the wild-type AAV1 or AAV2 rep gene (Figure 2C). This observation was surprising given the differential p40-promoter activities (Figure 2A). In contrast, swapping the zinc-finger domain sequences to the respective other AAV serotype resulted in significant differences of VP and Rep expression. The R2z1V1 construct showed a reduction of VP expression and a lower expression of Rep68/52 compared to Rep78/52 (Figure 2C). Vice versa, R1z2V1 showed an increase of VP expression and an approximately equal expression of Rep68/52 to Rep78/52 unlike the R1V1 construct. Thus, it is possible that these observations are linked, particularly since the VPs are translated from spliced p40 transcripts and mutations of the splice sites result in a strong reduction of VP expression (30, 31). In absence of nucleotide differences of the splice site in the AAV1 *rep* gene, the z-region, which comprises the majority of the intron sequence, has to be responsible for the inefficient splicing. Further research will be needed to determine whether the splice reduction is caused by the zinc-finger domain of the Rep proteins or by the differences of the DNA sequences. However, for this study the rescue of VP expression was achieved, and all subsequent variants contained the 3’end of the AAV2 *rep* gene.

### Rescue of AAV8 VP expression by correction of the AAV8 Rep DNA binding domain

The substitution of the 3’end of the *rep* gene to the sequences of AAV2 rescued AAV1 VP expression as shown above (Figure 2C) and AAV6 VP expression (data not shown). Thus, the same strategy was pursued in the case of AAV8 vector production using the AAV8 *rep* gene (Figure 3A). However, unlike the AAV1 and AAV6 constructs, no significant AAV8 VP expression was observed with the R8c2V8 construct (Figure 3B), comparable to the R8V8 construct as seen before (Figure 1B). Furthermore, the inability to express VP proteins was not just restricted to AAV8, as AAV1 and AAV6 VP expression is also inhibited when the AAV8 rep gene was utilized (data not shown). Vice-versa, AAV8 VP expression can be restored when using the AAV1 *rep* gene with the AAV2 3’ end (R1c2V8) to a similar level compared to the standard AAV2 *rep* system (Figure 3B). This observation was surprising given the 95% amino acid sequence identity between AAV1 Rep and AAV8 Rep (Table 1). In order to identify the region of the AAV8 *rep* gene responsible for this phenotype *rep* was sectioned into three additional parts (n, d, and h) and substituted to the corresponding AAV1 sequences (Figure 3A). Significant AAV8 VP expression was only achieved when the d-region was substituted for the AAV1 sequences (R8d1c2V8, Figure 3B) which is equivalent to the sequence encoding the DNA-binding domain. Further experiments showed that the lack of efficient AAV8 VP expression with the AAV8 rep gene can also be rescued by the addition of an AAV1 Rep expression construct (p5-Rep1) *in trans* (Figure 3C). Conversely, an AAV8 Rep expression construct (p5-Rep8) does not repress AAV8 VP expression when using the standard AAV2 Rep system or enhance VP expression in the R8c2V8 construct. These observations point to a defect of the AAV8 Rep protein, that is associated with the DNA binding domain. A sequence analysis of AAV8 to AAV1 Rep (and other AAV serotypes) in the affected region revealed two highly variable regions termed VR-A (residues 117-126, with 8 amino acid substitutions and 1 insertion) and VR-B (residues 137-143, with 5 amino acid substitutions and 1 insertion) (Figure 3D). However, when the sequences were analyzed on DNA sequence level only two nucleotide exchanges and 3 insertions were found for VR-A and one nucleotide exchange and 3 insertions for VR-B, respectively. The fact that the individual nucleotide insertions of AAV8 are spread out results in a shift of the reading frame relative to AAV1 and causes the high amino acid variability despite little differences at the DNA level. Since both VRs contain 3 nucleotide insertions the ribosomes shift back into the original reading frame during translation (Figure 3D). The occurrence of the 3 spaced-out nucleotide insertions is unlikely to happen naturally as they would need to be introduced simultaneously. Otherwise, the individual insertions would shift the reading frame and truncate the Rep proteins significantly. Thus, it is more likely that these insertions represent sequencing errors of the deposited AAV8 *rep* gene sequence. Therefore, the nucleotide sequences of VR-A and VR-B were corrected by removing the nucleotide insertions relative to AAV1 which resulted in a highly similar amino acid sequences with only a single amino acid substitution in VR-A and the identical amino acid sequence in VR-B (Figure 3D). Following transfection of these constructs the AAV8 VP expression was analyzed. While the correction of VR-A alone did not improve VP expression, the construct with the corrected VR-B showed significant VP expression (Figure 3E). This expression was further enhanced when both corrected regions were combined resulting in a similar VP expression compared to the R8d1c2V8 construct. The rescue of AAV8 VP expression can be explained based on a previously determined X-ray crystallography structure of AAV2 Rep bound to its binding element within the AAVS1 sequence (32). A model generated for the uncorrected AAV8 Rep protein based on the AAV2 Rep structure placed VR-A distance from, but VR-B directly in the major groove of the bound DNA (Figure 3F). While VR-A does not directly bind to the DNA, it could stabilize the binding of VR-B to the DNA and thus, further enhance VP expression. As the Rep proteins have been described as an activator of AAV transcription (33) it is possible that the defective AAV8 Rep proteins are unable to transactivate the p40 promoter (34) needed for efficient VP expression.

**Figure 3:**
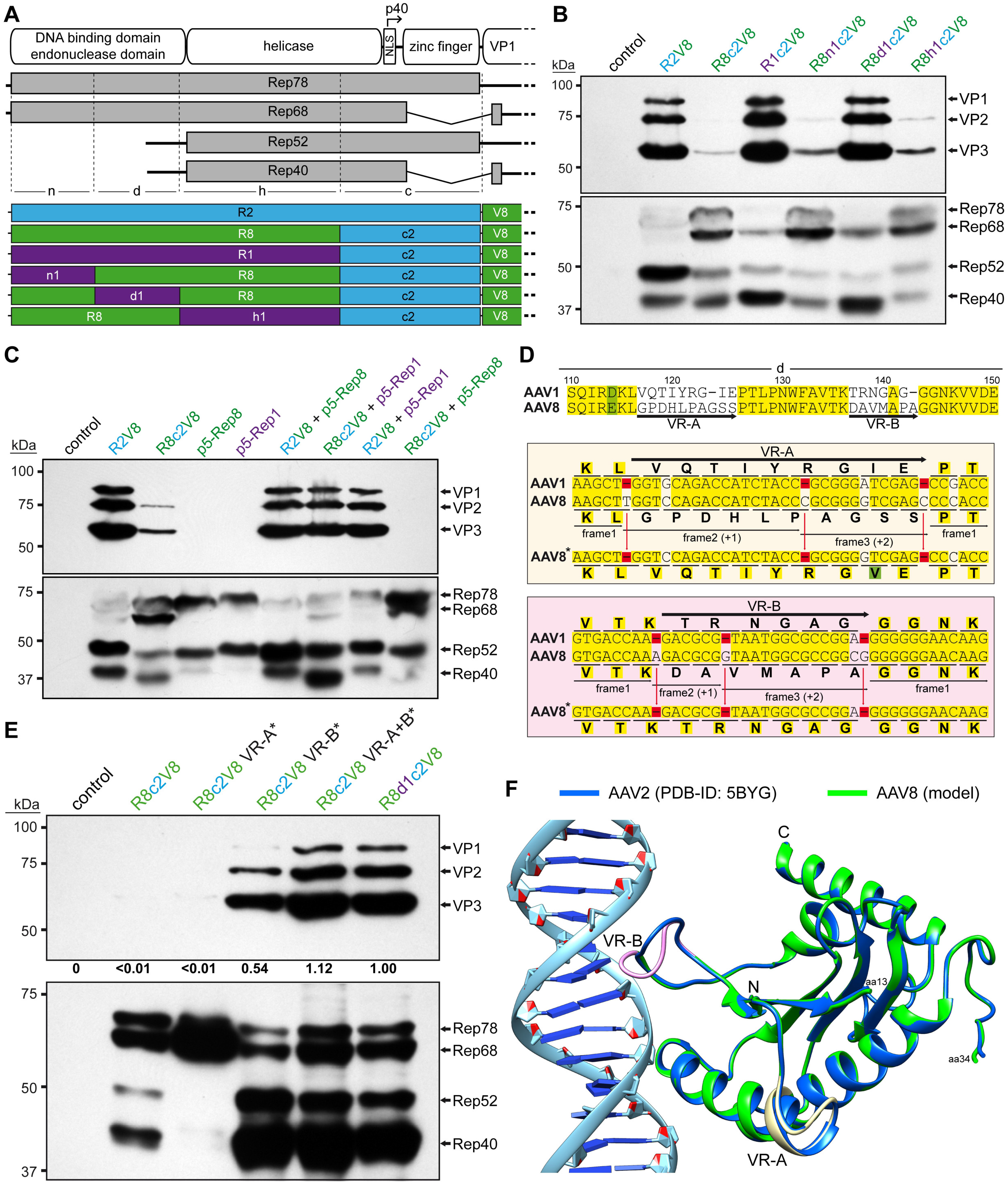
Rescuing VP expression utilizing the AAV8 *rep* gene. **A)** Schematic depiction of the Rep protein with its main domains. The approximate position of the p40 promoter in the *rep* gene is indicated. Below the p5 and p19 transcripts are shown with the translated regions for Rep78, Rep68, Rep52, and Rep40. Rep has been subdivided into a N-terminal domain (n), the DNA-binding domain (d), the helicase domain (h), and a C-terminal domain (c). The generated constructs containing different domains from AAV1, AAV2, and AAV8 are depicted below. Rep gene fragments derived from AAV1 are colored in purple, AAV2 in blue, and AAV8 in green. **B)** Western-blot analysis to determine Rep and VP expression of the constructs shown in A. Top blots were probed with MAb B1 and lower blots with MAb 1F. The individual VPs and Rep proteins are indicated. **C)** Analysis as in (B) in presence or absence of co-transfected Rep constructs. **D)** Amino acid sequence alignment of AAV1 and AAV8 in the DNA-binding domain (aa110 to aa150). Identical residues are highlighted in yellow and amino acid with similar properties in green. The two regions of significant amino acid difference termed VR-A and VR-B are indicated. Below an analysis of these regions are shown at nucleotide level with the encoded amino acids shown. Deletions in AAV1 relative to AAV8 are highlighted in red. The insertions in AAV8 potentially result in shifts to an alternative reading frame. Removal of these insertion (AAV8*) result in a similar amino acid sequence to AAV1. **E)** Analysis as in (B) utilizing the AAV8 Rep variants with the removed insertions. The intensity of the VP bands was quantified using ImageJ and normalized to R8d1c2V8. **F)** Structural analysis of the DNA-binding site for AAV2-Rep (blue) and a superposed AAV8-Rep model (green) to AAVS1 dsDNA. The position of VR-A and VR-B is indicated.

### AAV1/2 Rep hybrids can enhance vector genome packaging efficiency into AAV1 capsids

Initial analyses of lysates showed that vector genome titers were comparable to the standard AAV2 rep system (R2V1) when the ATG-R1V1 construct was used despite displaying lower VP expression (Figure 4A). Following the rescue of the VP expression with ATG-R1c2V1, the genome titer was higher than R2V1 but at a similar VP expression level. However, this enhancement was only observed for the ATG start codon constructs while the ACG constructs showed very low genome titers (Figure 4A).

**Figure 4:**
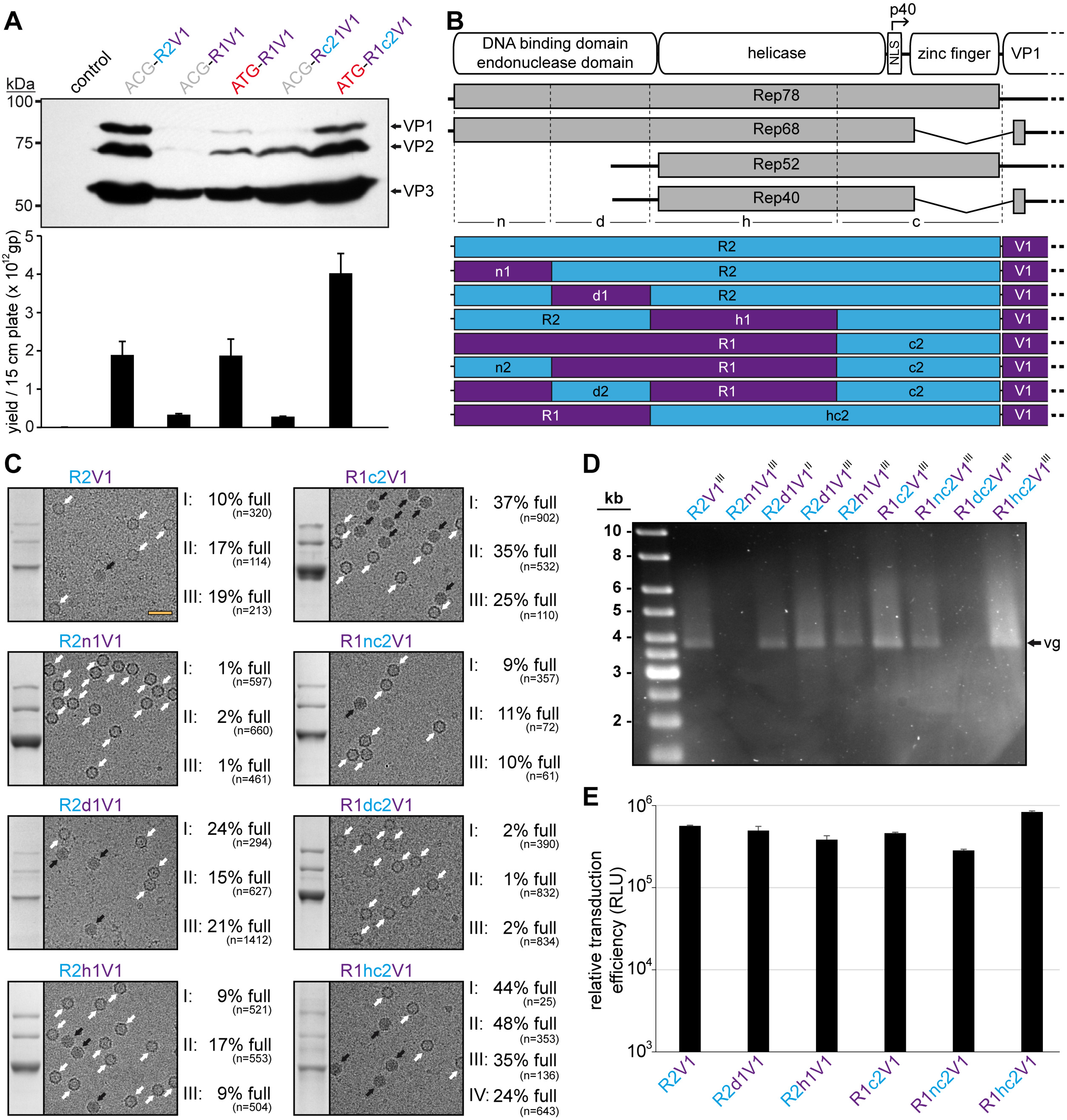
AAV1/2 Rep hybrids improve AAV1 genome packaging efficiency. **A)** Analysis of VP expression by Western-blot and the AAV1 vector yield by qPCR following transfection of the constructs in HEK293 cells. The Western blot was probed with MAb B1. The individual VPs and Rep proteins are indicated. **B)** Schematic depiction of the Rep protein with its main domains. The approximate position of the p40 promoter in the *rep* gene is indicated. Below the p5 and p19 transcripts are shown with the translated regions for Rep78, Rep68, Rep52, and Rep40. Rep has been subdivided into a N-terminal domain (n), the DNA-binding domain (d), the helicase domain (h), and a C-terminal domain (c). The generated constructs containing different domains from AAV1 and AAV2 are depicted below. Rep gene fragments derived from AAV1 are colored in purple and AAV2 in blue. **C)** Analysis of the AVB-purified AAV1 vector preparations. Sections of SDS-PAGEs containing VP1, VP2, and VP3 and representative example cryo-EM micrographs are shown for each Rep hybrid. White arrows point to empty capsids (light appearance) and black arrows to full capsids (dark appearance). The determined percentage of full capsids of at least three independently produced and purified AAV1 vector preparations are displayed with the total particle count of all micrographs collected for the individual sample. Scale bar (shown in R2V1 micrograph): 50 nm. **D)** Alkaline gel electrophoresis of the AAV1 vector preparations. Capsid amount loaded is the same for all samples based on the ELISA titer. The size of the packaged vector genome (vg) is ~3.9 kb. **E)** Analysis of the transduction efficiency of the AAV1 vectors produced with different Rep hybrids in HEK 293 cells.

Thus, the genome packaging efficiency of the Rep hybrids into AAV1 capsids was further investigated. For this purpose, the *rep* genes of AAV1 and AAV2 were divided into 3 additional regions (n, d, and h) similar as described for AAV8 above to identify which region provides a potential benefit for genome packaging (Figure 4B). For these regions eight possible permutations between AAV1 and AAV2 were generated (R2, R2n1, R2d1, R2h1, R1c2, R1nc2, R1dc2, and R1hc2) and utilized for AAV1 vector production. As the packaging efficiency was expected to vary slightly between different vector preparations three independent vector preparations for each Rep variant were produced. The individual AAV1 vector preparations were purified also independently by AVB affinity chromatography that indiscriminately binds empty and full capsids (35). Subsequently, each of the AAV1 vector preparations were analyzed by cryo-electron microscopy (cryo-EM) and the percentage of full capsids determined (Figure 4C). For the standard AAV2 Rep production system (R2V1) the percentage of full capsids in the three vector preparations ranged from ~10-20% with a mean average of 15% (Table 2). In contrast, for AAV1 Rep with the AAV2 C-terminus (R1c2V1) the percentage of full capsids in the three vector preparations is approximately doubled and ranged from 25-37% with a mean average of 32%. Swaps of the N-termini based on these constructs to the corresponding other AAV serotype was detrimental to both constructs. While the AAV1 N-terminus in AAV2 Rep (R2n1V1) resulted in a vast majority of empty capsids (% full: 1-2%, average: ~1%), the AAV2 N-terminus in AAV1 (R1nc2V1) reduced the percentage of full capsids to 9-11% with a mean average of 10% (Figure 4C, Table 2). Similarly, the AAV2 DNA-binding domain inserted into AAV1 Rep (R1dc2V1) also resulted in largely empty capsids (% full: 1-2%, average: ~2%). These results could indicate that the AAV2 DNA binding domain is not compatible with the AAV1 N-terminus which are expressed by the R2n1V1 and R1dc2V1 constructs. Vice versa, the AAV1 DNA-binding domain is supported in AAV2 Rep and slightly enhanced packaging efficiency (% full: 15-24%, average: ~20%). Swapping the AAV1 helicase domain into AAV2 (R2h1V1) did not affect the packaging efficiency significantly (% full: 9-17%, average: ~12%) compared to R2V1. Lastly, the Rep hybrid with the best overall packaging efficiency contained the AAV2 helicase within AAV1 Rep (R1hc2V1) with a percentage of full capsids in the four vector preparations ranging from 24-48% with a mean average of 38% (Figure 4D, Table 2). The approximate full-empty ratios for the different Rep hybrids were also confirmed by qPCR when compared to the capsid titer determined by ELISA (data not shown). The packaged genomes were also visualized by performing an alkaline gel electrophoresis loading equal amounts of capsids (Figure 4D). Thus, vector preparations with a higher percentage of full capsids will appear brighter such as for R1hc2V1 whereas no bands are seen for largely empty capsids (R2n1V1 and R1dc2V1). Regardless of the utilized Rep hybrid during vector production, no significant differences of the transduction efficiency of the purified AAV vectors were observed (Figure 4E). This was expected as the Rep proteins are not part of the final purified AAV vector preparations.

**Table 2:** Summary of the quantification of the full capsids for AAV1

| AAV serotype<br>(capsid) | Utilized<br>Rep variant | biological<br>replicate | percentage<br>"full" capsids | mean |
| --- | --- | --- | --- | --- |
| AAV1 | R2 | I | 10% | 15% |
|  |  | II | 17% |  |
|  |  | III | 19% |  |
|  | R2n1 | I | 1% | 1% |
|  |  | II | 2% |  |
|  |  | III | 1% |  |
|  | R2d1 | I | 24% | 20% |
|  |  | II | 15% |  |
|  |  | III | 21% |  |
|  | R2h1 | I | 9% | 12% |
|  |  | II | 17% |  |
|  |  | III | 9% |  |
|  | R1c2 | I | 37% | 32% |
|  |  | II | 35% |  |
|  |  | III | 25% |  |
|  | R1nc2 | I | 9% | 10% |
|  |  | II | 11% |  |
|  |  | III | 10% |  |
|  | R1dc2 | I | 2% | 2% |
|  |  | II | 1% |  |
|  |  | III | 2% |  |
|  | R1hc2 | I | 44% | 38% |
|  |  | II | 48% |  |
|  |  | III | 35% |  |
|  |  | IV | 24% |  |

### The Rep hybrids also enhance vector genome packaging efficiency for other AAV serotypes

A similar screen for the genome packaging efficiency, as done for AAV1 (Figure 4A), was conducted for AAV6 utilizing AAV6 hybrids with the AAV2 C-terminus. However, the AAV1 and AAV8 *rep* hybrids cloned upstream of the AAV6 *cap* gene appeared to exceed the packaging efficiency over the AAV6 Rep hybrids (data not shown). Thus, the AAV1 and AAV8 Rep hybrids were utilized for AAV6 production and compared to the standard AAV2 Rep system (Figure 5A). Similar to AAV1 the three independent AAV6 vector preparations were purified by AVB-affinity chromatography and the percentage of full capsids determined by cryo-EM imaging. For the standard AAV2 Rep system (R2V6) the percentage of full capsids ranged from 12-20% (average 17%) (Table 3). In contrast, the percentage of full capsids is increased for each of the other tested Rep hybrids ranging from 20-39% (average 28%) for R2d1V6, 37-46% (average 43%) for R1hc2V6, and 32-38% (average 36%) for R8d1c2V6 (Figure 5A, Table 3). Due to the high sequence similarity of the AAV1 and AAV6 capsids (36) it was not surprising that the Rep hybrids improved packaging efficiency similarly.

**Figure 5:**
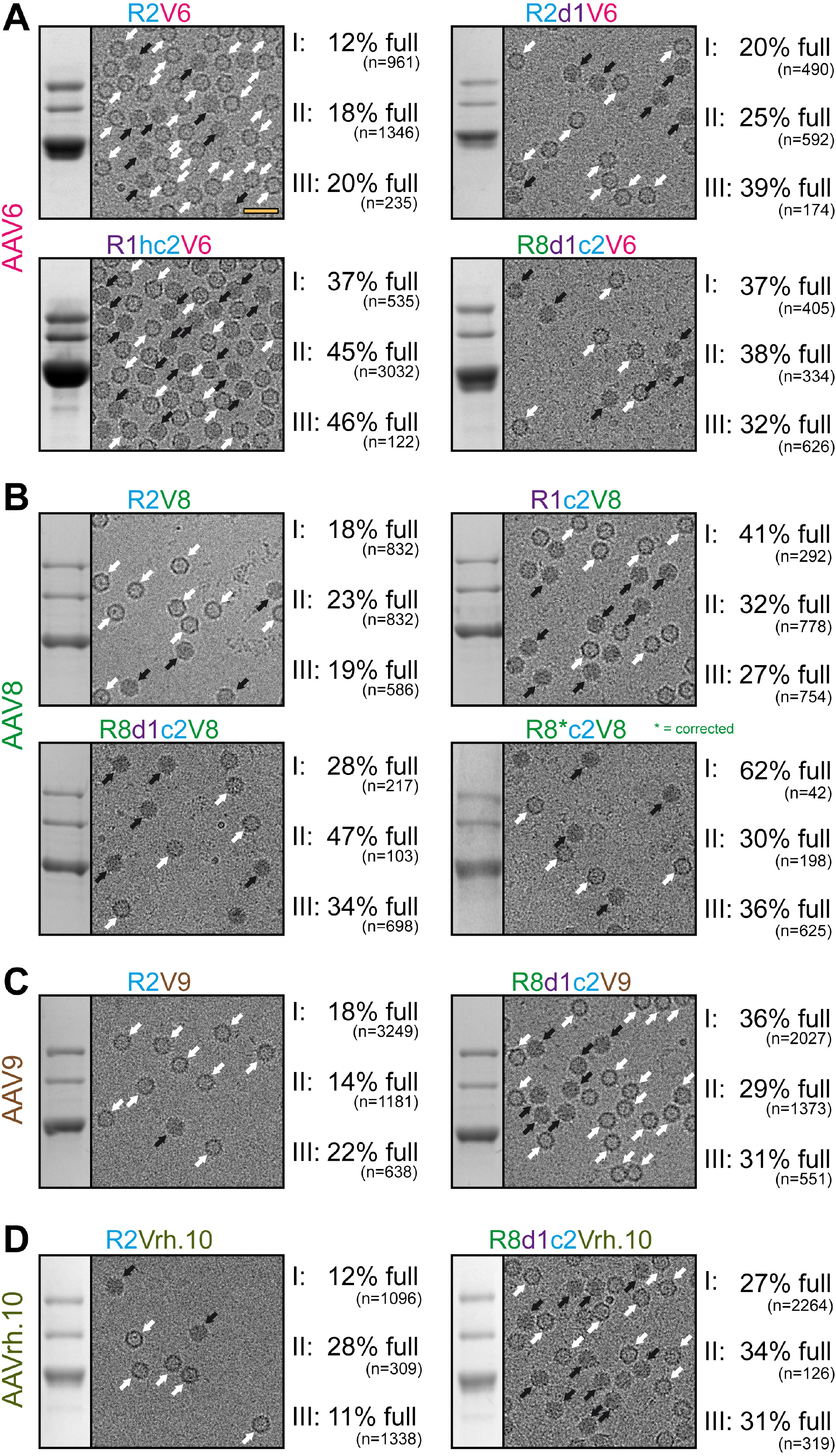
The Rep hybrids also enhance packaging efficiency for other AAV serotypes. **A)** Analysis of the AVB-purified AAV6 vector preparations. Sections of SDS-PAGEs containing VP1, VP2, and VP3 and representative example cryo-EM micrographs are shown for each Rep variant. White arrows point to empty capsids (light appearance) and black arrows to full capsids (dark appearance). The determined percentage of full capsids of three independently produced and purified AAV6 vector preparations are displayed with the total particle count of all micrographs collected for the individual sample. Scale bar (shown in R2V6 micrograph): 50 nm. **B)** Analysis as in (A) for AAV8-Capture select affinity ligand purified AAV8 vectors. **C)** Analysis as in (A) for AAV9-Capture select affinity ligand purified AAV9 vectors. **D)** Analysis as in (A) for AVB-purified AAVrh.10 vectors.

**Table 3:** Summary of the quantification of the full capsids for AAV6, 8, 9, and rh.10

| AAV serotype<br>(capsid) | Utilized<br>Rep variant | biological<br>replicate | percentage<br>"full" capsids | mean |
| --- | --- | --- | --- | --- |
| AAV6 | R2 | I | 12% | 17% |
|  |  | II | 18% |  |
|  |  | III | 20% |  |
|  | R1hc2 | I | 37% | 43% |
|  |  | II | 45% |  |
|  |  | III | 46% |  |
|  | R2d1 | I | 20% | 28% |
|  |  | II | 25% |  |
|  |  | III | 39% |  |
|  | R8d1c2 | I | 37% | 36% |
|  |  | II | 38% |  |
|  |  | III | 32% |  |
| AAV8 | R2 | I | 18% | 20% |
|  |  | II | 23% |  |
|  |  | III | 19% |  |
|  | R8d1c2 | I | 28% | 36% |
|  |  | II | 47% |  |
|  |  | III | 34% |  |
|  | R1c2 | I | 41% | 33% |
|  |  | II | 32% |  |
|  |  | III | 27% |  |
|  | R8*c2 | I | 62% | 43% |
|  |  | II | 30% |  |
|  |  | III | 36% |  |
| AAV9 | R2 | I | 18% | 18% |
|  |  | II | 14% |  |
|  |  | III | 22% |  |
|  | R8d1c2 | I | 36% | 32% |
|  |  | II | 29% |  |
|  |  | III | 31% |  |
| AAVrh.10 | R2 | I | 12% | 17% |
|  |  | II | 28% |  |
|  |  | III | 11% |  |
|  | R8d1c2 | I | 27% | 31% |
|  |  | II | 34% |  |
|  |  | III | 31% |  |

For AAV8 vectors produced with AAV2 *rep* and the percentage of full capsids ranged from 18-23% (average 20%) (Figure 5B, Table 3). The utilization of the Rep hybrids that also rescued AAV8 VP expression (Figure 3) improved the percentage of genome-containing capsids to 27-41% (average 33%) for R1c2V8, 28-47% (average 36%) for R8d1c2V8, and 30-62% (average 43%) for the corrected R8c2V8. Following the correction of the AAV8 DNA binding domain (Figure 3D) the resulting Rep protein of the R8c2V8 construct varies only in two amino acids from the R8d1c2 Rep hybrid. Thus, a similar level of enhancement of genome packaging efficiency was expected.

For AAV9 and AAVrh.10 the *rep* genes or Rep protein sequences have not been isolated or deposited (21). Alternatively, the most promising Rep hybrids for AAV1, AAV6, and AAV8 were cloned upstream of the AAV9 or AAVrh.10 *cap* gene and the yield of the constructs by qPCR compared to the VP expression intensity by Western-blot (data not shown). For both AAVs the R8d1c2 Rep hybrid came ahead of the standard AAV2 Rep and three independent AAV preparations were generated with either Rep variant as with the previous AAV serotypes for detailed analysis. For AAV9 the percentage of full capsids ranged from 14-22% (average: 18%) with R2V9 whereas genome-containing capsids were increased to 29-36% (average: 32%) with R8d1c2V9 (Figure 5C). For AAVrh.10 the results were similar with 11-28% (average 17%) full capsids with R2Vrh.10 and 27-34% (average 31%) full capsids with R8d1c2Vrh.10 (Figure 5D, Table 3).

These results indicate that the new Rep hybrids may improve packaging regardless of which capsids are utilized as the amino acid sequence identity of these capsids vary from 82-99% (Table 1). In order to confirm this conclusion, the effects of the Rep hybrids were also analyzed on AAV2 vector production. As expected, the overall yield of capsids with the different Rep hybrids were comparable (Figure 6A) but the vector genome cassettes were packaged equally (Figure 6B). As previously mentioned, packaging of vector genomes into AAV2 capsids is generally more efficient (23). This was also observed using cryo-EM imaging with the R2V2 construct generating ~47% of full capsids (Figure 6C). In contrast, the percentage of full capsids for the other AAV serotypes ranged from 12-20% when the AAV2 *rep* gene was used (Figure 4C, Figure 5). With the different Rep hybrids, the percentage of full AAV2 capsids ranged between 32 – 65% (Figure 6C) which corresponds to a 0.7-1.4-fold decrease or increase relative to the wtAAV2 Rep construct, respectively. As the portion of genome-containing capsids was high for AAV2 to start with, a further improvement of packaging was not expected. On the other hand, utilizing the Rep hybrids such as R1c2 or R8d1c2 for AAV2 vector production the percentage of full AAV2 capsids did not drop significantly either. This questioned the initial hypothesis that the Rep proteins of an AAV serotype might be adapted to its own capsid. Thus, there has to be an alternative explanation for the higher percentage of full capsids when using these Rep hybrids. The Rep hybrids that allowed the best packaging efficiency contained the ~240 N-terminal amino acids of either AAV1 or AAV8 Rep78/68 which contains the DNA-binding and endonuclease domain of the Rep proteins (16). This region is indispensable for genome replication (18, 37, 38). Thus, it is possible that the Rep hybrids replicate the vector genomes to higher copy numbers prior to their packaging. However, the copy number of vector genomes determined by qPCR after lysis of transfected cells without DNase treatment showed similar level of replicated genomes for all Rep variants (data not shown). The small Rep proteins (Rep52/40) have been suggested to be responsible for the encapsidation of the genomes into the capsids (18). However, the small Rep proteins are largely identical between Rep2 and the R1hc2 variant (Figure 4D) except for a single amino acid substitution (K234R). Thus, it is likely that the large Rep proteins are also involved in the encapsidation process with the Rep hybrids improving genome packaging by a currently unknown mechanism.

**Figure 6:**
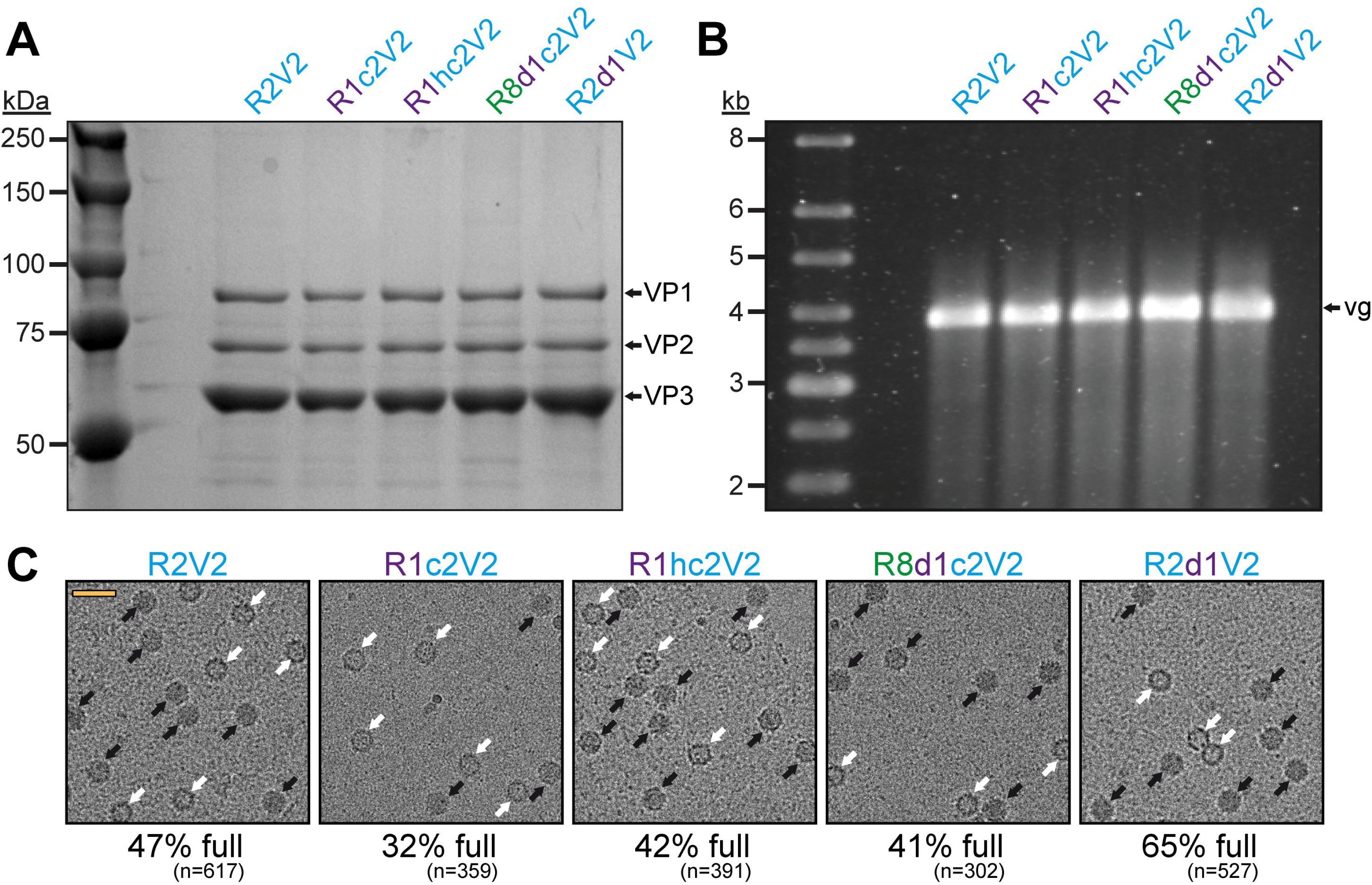
The effect of the Rep hybrids on genome encapsidation of AAV2. **A)** SDS-PAGE of the AVB-purified AAV2 vector preparations produced with different Rep variants. VP1, VP2, and VP3 are indicated. **B)** Alkaline gel electrophoresis of the AAV2 vector preparations. Capsid amount loaded is the same for all samples. The size of the packaged vector genome (vg) is ~3.9 kb. **C)** Representative example cryo-EM micrographs are shown for each Rep variant. White arrows point to empty capsids (light appearance) and black arrows to full capsids (dark appearance). The determined percentage of full capsids of the vector preparations are displayed with the total particle count of all micrographs collected for the individual sample. Scale bar (shown in R2V2 micrograph): 50 nm.

## Conclusions

The vast majority of AAV production systems utilize AAV2 *rep* for vector manufacturing. The few exceptions reported previously include the utilization of the *rep* gene of AAV3 (39) and AAV4 (22) for production. In both cases the vector yield was described to be higher, but no further investigation was conducted. In this study the *rep* genes of different AAV serotypes were analyzed and while they were not directly usable “as is”, they were modified to generate hybrids supporting high capsid expression and improved packaging efficiency. The fact that the Rep hybrids enhanced packaging efficiency across multiple, widely used AAV serotypes in gene therapy trials indicates that they can be possibly used for almost all AAV capsids to improve overall vector yield and increase of the percentage of genome-containing capsids. As the *rep* gene is often provided *in trans* for most AAV production system the new hybrid *rep* genes can easily replace the AAV2 *rep* constructs. Furthermore, during AAV vector purification the Rep proteins are removed, making the final product indistinguishable to the AAV2 *rep* system except for the higher content of genome-containing capsids. While the exact mechanism of the enhancement of packaging remains unclear further research on the Rep proteins is needed in general. It will be also interesting to see if the Rep hybrids also improve packaging in insect cells for large scale AAV production setups that often suffer from high empty-to-full capsid ratios (40). Empty capsids are generally not desired in AAV vector preparations as they do not provide any curative benefit for the treatment of the targeted disease and can reduce overall transduction efficiencies (41). Some purification protocols do not actively remove empty capsid which could be problematic when administered to a patient with a given genome-containing vector dose as the empty particle titer can be 5 to 10-fold higher. These empty capsids can then potentially elicit additional immune responses *in vivo* gene therapies.

## Materials and Methods

### Plasmids and cloning

The *rep* genes of AAV1, AAV6, and AAV8 were synthesized by GenArt (Thermo Fisher) based on the deposited AAV serotype genomes; accession numbers: AF063497 (AAV1), AF043303 (AAV2), and AF513852 (AAV8). These rep genes were inserted into plasmids containing the AAV2 *rep* gene and either the AAV1, 6, or 8 *cap* gene (R2V1, R2V6, or R2V8) by replacing the AAV2 *rep* gene to generate R1V1, R6V6, or R8V8, respectively. Site-directed mutagenesis was utilized by as previously described (42) to mutate the Rep78 start codon or to correct the AAV8 *rep* gene. In order to generate Rep hybrid proteins, the *rep* gene was subdivided into four regions and cloned using restriction sites that are conserved in equivalent positions of the AAV1, 2, 6, and 8 rep genes. Briefly, the N-terminal region (n) extends to *Nco*I (nt position 305 of the AAV1 *rep* gene, amino acid position 102 of AAV1 Rep), the DNA-binding domain (d) to *BamH*I (nt position 725, amino acid position 242), the helicase domain (h) to *Sal*I (nt position 1108, amino acid position 370) and the C-terminal region consists all the sequences past *Sal*I. In absence of usable restrictions site Gibson assembly was performed using the NEB Gibson assembly master mix according to the manufacturer instructions to further subdivide the C-terminal region at nt position 1590, amino acid position 530 (AAV1 numbering), termed y and z region. To determine the p40 promoter activity the nts 1442-1878 (AAV1), nts 1428-1887 (AAV2), and nts 1340-1776 (AAV8) from the accession numbers listed above were cloned upstream of a luciferase gene.

### Cell culture

HEK293 cells were maintained in Dulbecco’s Modified Eagle Medium (DMEM) supplemented with 10% heat-inactivated fetal calf serum and 100 U of penicillin/ml and 100 μg of streptomycin at 37°C in 5% CO_2_.

### AAV production and purification

Recombinant AAV vectors, with a packaged luciferase gene, were produced by triple transfection of HEK293 cells, utilizing pTR-UF3-Luciferase, pHelper (Stratagene), and a *rep*-*cap* plasmid containing either the wild-type or hybrid *rep* genes. The transfected cells were harvested 72 h post transfection as previously described (42). The cleared lysates containing AAV capsids were purified by AVB Sepharose (Thermo Fisher) in the case of AAV1, 2, and 6 and by POROS CaptureSelect™ AAV8 (Thermo Fisher) affinity chromatography as previously described (35).

### SDS-PAGE and Western-blot analysis

The purity of the AAV preparations were confirmed by sodium dodecyl sulfate polyacrylamide gel electrophoresis (SDS-PAGE). For this purpose, the samples were incubated with 6× Laemmle Sample Buffer (Bio-Rad) with 10% β-mercaptoethanol and boiled for 5 min at 100°C. The denatured proteins were applied to a 10% polyacrylamide gel and ran at 120 V. The gel was washed three times with distilled water (diH2O) and stained with GelCode Blue Protein Safe stain (Invitrogen). In order to confirm and evaluate Rep and Cap expression from the new Rep hybrid plasmids Western-blot analyses were performed. For this purpose, the proteins were transferred to a nitrocellulose membrane following SDS-PAGE by electroblotting. The membrane was blocked in 6% milk in 1xPBS and probed with hybridoma supernatants containing MAb B1, detecting VP1, VP2, and VP3, (43) or MAb 1F, detecting Rep78, Rep68, Rep52, and Rep40 (29). Following the incubation with a secondary antibody with a linked horse radish peroxidase the proteins were visualized by applying Immobilon™ Chemiluminescent Substrate (Millipore) and detection on an X-ray film.

#### Quantification of AAV vectors

Aliquots from the AAV vector preparations were digested with Proteinase K to release the AAV vector genomes from the capsids. To this end, the samples were incubated in buffer containing 10 mM Tris pH 8, 10 mM EDTA, 1% SDS for 2 h at 56°C. The released DNA was purified utilizing the PureLink PCR Purification Kit (Thermo Fisher). The copy number of vector genome DNAs were determined by quantitative PCR using iQ™ SYBR^®^ Green Supermix (BioRad, Hercules, CA). Primers specific for the luciferase gene of the vector genome were used (Forward primer 5’-GCAAAACGCTTCCATCTTCC-3’ and reverse primer 5’-AGATCCACAACCTTCGCTTC -3’).

#### AAV capsid ELISA

For the quantification of the physical capsid titer AAV Titration ELISA (Progen) were utilized. All the steps were done according to the protocol provided by the manufacturer in triplicate. The colorimetric assay was analyzed by a Synergy HT plate reader (BioTek).

#### Alkaline gel electrophoresis

For the alkaline gel electrophoresis, a 0.8 % agarose gel in 1xTAE buffer (40 mM Tris, 20 mM acetic acid, and 1 mM EDTA) was utilized. Following solidification, the gel was equilibrated in 1x denaturing buffer (0.5 M NaOH, 50 mM EDTA) for 4 h. Prior to the loading, the samples were mixed with denaturing loading dye (final concentration: 1x Ficoll loading buffer, 1x denaturing buffer, 10% SDS). The agarose gel was run at low voltage overnight at 4°C. After the run the gel was washed and neutralized in 1xTAE for 30 min and subsequently stained in a 0.02% SyBr-Gold solution in 1xTAE. The gel was imaged under UV light using a BioRad GelDoc system.

### Analysis of transduction efficiency and promoter activity

Purified AAV vectors with a packaged luciferase expression cassette were used to infect HEK-293 cells at a 10^5^ MOI (multiplicity of infection). After 48 h, cells were lysed and luciferase activity assayed using a luciferase assay kit (Promega) as described in the manufacturer’s protocol. Luminescence was measured on a Synergy HT plate reader (BioTek). In order to determine the p40 promoter activity in HEK293 cells the AAV-p40-luciferase constructs were transfected and the cells incubated for 24 hours. The subsequent steps were performed identically as described above for the analysis of transduction efficiency.

#### Cryo-Electron microscopy imaging

For each of the purified AAV capsids, 3.5 μL was applied to a glow-discharged Quantifoil copper grid with 2 nm continuous carbon support over holes (Quantifoil R 2/4 400 mesh), blotted, and vitrified using a Vitrobot Mark 4 (FEI) at 95% humidity and 4°C. Images were collected using an FEI Tecnai G2 F20-TWIN microscope (FEI) operated under low-dose conditions (200 kV, ~20e^−^/Å^2^) on a GatanUltraScan 4000 CCD camera (Gatan). The number of empty and full capsids in these images were counted manually.

## Acknowledgments

The authors thank the UF-ICBR Electron microscopy core for access to electron microscopes utilized for cryo-electron micrograph screening.

